## Supplementary Materials for "Transsaccadic working memory in healthy ageing and neurodegenerative disease"

### Larger saccades disrupt memory more

To investigate whether larger saccades disrupt spatial memory more, we conducted a multiple linear regression analysis. This model predicted location error size based on four factors: horizontal and vertical saccade direction (left/right and up/down, respectively), and horizontal and vertical distance between stimulus frames (***Figure 2 – supplementary figure 1A***). While the horizontal direction coefficient showed marginal significance (t(41) = -1.94, p = 0.059), only the vertical distance coefficient was robustly significant (t(41) = 5.73, p < 0.0001). This effect was consistent across both age groups. These results indicate that spatial memory impairment increases with greater distance between the item and the subsequent gaze target, in particular the vertical axis, though regardless of which direction the eye is moving to.

While the direction of gaze after item representation does not appear to differentially affect memory, we further investigated, amongst transsaccadic trials where two saccades were made, whether crossing saccades (e.g., a rightward saccade followed by a leftward saccade) might reduce spatial memory precision compared to consecutive saccades in a single direction. A repeated measures ANOVA revealed no interaction between horizontal direction (leftward vs. rightward vs. horizontal crossing saccades) and age group (Young vs Elderly) (***Figure 2 – supplementary figure 1B***, F(1.3,49.8)=0.04, p=0.89, partial η²=0.001). Although older participants had significantly larger error overall (main effect of age group: F(1, 39)= 9.00, p=0.005, partial η²=0.19), there was no main effect of direction (F(1.3,49.8)=0.64, p=0.47, partial η²=0.02). Similarly, a repeated measures ANOVA for vertical saccade direction (upward vs. downward vs. vertical crossing) showed no interaction with age group (***Figure 2 – supplementary figure 1C***, F(1.7,62.8)=0.29, p=0.71, partial η²=0.008). Again, older participants demonstrated greater overall error (main effect of age group: F(1, 38) = 15.00, p < 0.001, partial η² = 0.28), but there was no main effect of vertical direction (F(1.7, 62.8) = 1.41, p = 0.25, partial η² = 0.04). This is expected as these participants are healthy and would not be expected to have a systematic lateralised deficit.

### Exploration of ROCF Drawing Dynamics and the relation with the LOCUS

We conducted a more in-depth analysis of ROCF drawing behaviour, excluding two elderly healthy participants due to accidental loss of their real-time recordings (N=40). First, we examined the time taken to complete the ROCF copy. Young participants completed the copy drawing in approximately 2 minutes (mean = 134.50 s, SD = 55.24), while older participants took significantly longer (mean = 221.13 s, SD = 104.77; t(38) = -3.32, p < 0.001). However, the duration did not correlate with copy performance (rho = 0.18, p = 0.27, Fisher’s z = 0.18, BF_10_ = 0.20) nor with saccade cost in the LOCUS task (rho = -0.14, p = 0.38, Fisher’s z = -0.14, BF_10_ = 0.20).

There was a robust positive correlation between retinotopic working memory deficit (no-saccade location error) and ROCF copy duration (rho = 0.50, p = 0.001, Fisher’s z = 0.55, BF_10_ = 7.93). A mediation analysis with retinotopic working memory as the predictor and ROCF copy duration as the outcome, considering total distance drawn, path drawing speed, and mean waiting time between paths as potential mediators, revealed a significant indirect effect only through mean waiting time between paths (i.e., the duration between completing the last path and starting the next path) (z = 3.33, *p* < 0.001, standardised estimate = 0.34 with confidence interval [0.17 0.64]). There was no direct effect of retinotopic working memory on copy duration (z = 0.07, *p* = 0.95, standardised estimate = 0.002 with confidence interval [-0.07 0.06]). This suggests that greater deficits in retinotopic working memory led to longer pauses between drawing strokes, potentially reflecting increased encoding time or decision-making difficulties. Eye-tracking data were not collected during the ROCF copy task, precluding further investigation of gaze patterns during drawing.
